# Evaluating State-Transition Models in Short Resting-State fMRI with Continuous-Dynamics Controls

**DOI:** 10.64898/2026.11.04.763846

**Authors:** Aleksandr Volkov, Yury Maximov, Tamoghna Chattopadhyay, Paul M. Thompson, Ildar Gabitov

## Abstract

Hidden Markov models (HMMs) are widely used to describe changes in resting-state functional MRI as transitions between a finite set of inferred brain states. However, continuously varying brain activity can produce similar apparent transitions, raising questions about their biological validity. We tested whether discrete and continuous brain dynamics could be distinguished by predicting subsequent fMRI measurements, and if temporal filtering affects this distinction. We compared hidden Markov models (HMMs) with vector autoregressive (VAR) models, which predict signals from preceding measurements without assuming discrete states. Each time point contained signals from the seven Yeo functional networks. Both models used the same preceding observations to predict the next, with accuracy measured by the log probability density assigned to the observed signals. Across 864 simulated conditions, HMMs performed better in 139/144 unfiltered discrete-state cases, whereas VAR performed better in 67/72 continuous cases. Changing only the filter during testing increased the number of continuous cases favoring HMMs from 2/72 with fixed filtering to 68/72, despite unchanged underlying dynamics. We also analyzed resting-state fMRI from 270 ADNI participants (90 cognitively normal, 90 with mild cognitive impairment, and 90 with dementia). VAR outperformed HMM in 160/184 datasets with at least 180 time points and 60/86 shorter recordings. In simulations, HMMs inferred directional state transitions even from smoothly evolving activity without discrete switching. Thus, apparent fMRI state transitions do not necessarily indicate discrete underlying dynamics. Their interpretation requires comparison with continuous models and attention to temporal filtering.

## 1 Introduction

Large-scale functional brain networks change over time [1, 2], offering insights into brain organization and its disruption in neurological and psychiatric disorders. Hidden Markov models (HMMs) represent brain activity using a finite set of latent states, each associated with a probability distribution of observed activity, and estimate transitions between states [3]. HMM studies have identified transient networks in electrophysiological recordings and hierarchical temporal organization in fMRI [4, 5]. Autoregressive and Gaussian-linear extensions model more complex relationships within states [6, 7]. However, fitted transitions need not reflect discrete neural changes [7]. In a short recording, a continuous trajectory may pass repeatedly through regions assigned to different states, producing apparent directional transitions without switching between distinct physiological configurations.

Liégeois et al. [8] showed that a first-order autoregressive model reproduced fluctuations in sliding-window functional connectivity more accurately than an HMM, illustrating how continuous models can explain apparent changes in network organization. Ahrends et al. [9] found that HMM estimates depend on the number of observations relative to model parameters and the choice of parcellation. Leonardi and Van De Ville [10] showed that filtering and window length influence estimates of connectivity fluctuations. These studies raise the question: *how reliably can inferred state transitions be distinguished from temporal structure arising from continuous dynamics or measurement choices?*

To investigate this, we compared HMMs with vector autoregressive (VAR) models, which predict each network signal from preceding measurements without discrete latent states. Both models used the same observed history to predict subsequent time points, and we compare their predictive log densities. We first tested them on simulated recordings generated by discrete state transitions or continuous shared fluctuations and rotational dynamics. With the underlying process known, we could assess whether each model family is favored under the expected conditions [11]. We then changed the observation filter during testing, keeping the underlying dynamics and fitted models fixed, to determine whether model preference changes. Separately, we measured the directional imbalance of fitted state transitions to test whether it distinguishes discrete switching from continuous motion. Finally, we applied this comparison to resting-state fMRI from 270 ADNI participants. Our goal was to assess what evidence HMM transitions provide for discrete dynamics and how that evidence depends on scan length and signal processing.

## 2 Data and Methods

### 2.1 Participants

We analyzed resting-state fMRI data from the Alzheimer’s Disease Neuroimaging Initiative (ADNI), comprising 4,347 BOLD time series from 4,257 visits by 1,764 participants [12]. We required a clinical diagnosis within 90 days of scanning, and sufficient fMRI data for analysis. This left 3,810 visits from 1,698 participants; 907 had at least two visits. Additional requirements for a T1-weighted scan from the same study and recorded sex information reduced this to 903 participants. To balance diagnostic groups and reduce differences in acquisition and demographic characteristics, we matched each of the 90 eligible participants with dementia to one cognitively normal (CN) participant and one with mild cognitive impairment (MCI). Matching also required the same number of eligible visits. The resulting sample contained 270 participants and 732 visits. For the present analysis, we used the first eligible scan from each participant, giving 270 independent recordings.

To reduce the complexity of modeling short fMRI recordings, we averaged the time series from 400 cortical regions in the Schaefer atlas [13] within seven predefined functional networks [14], giving one signal per network. We used the data before regression of 24 head-motion terms (M24), retaining motion-related variation in the measured signals. Because our aim was to compare models of brain activity rather than predict clinical diagnosis, diagnostic information was not used in fitting or selecting models. Both models were evaluated using the same preceding fMRI time points to predict subsequent measurements within each recording. To examine whether recording length affected performance, we analyzed 184 scans containing at least 180 time points and 86 shorter scans.

### 2.2 Discrete and Continuous Generators

To test whether HMMs can distinguish discrete state transitions from continuous dynamics, we generated synthetic recordings using four models with known underlying processes. Two models switched between three discrete states, while the other two produced continuously varying signals without discrete switching. All four generated a seven-dimensional time series, matching the seven brain networks used in the human fMRI analysis.

For the discrete models, we compared a process with a preferred direction of transitions between states (directed cycle) with one in which transitions were equally likely in either direction (reversible chain). Their transition probability matrices were

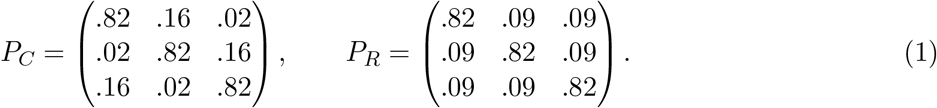

Both models had the same probability of remaining in each state (0.82), equal long-term state occupancy, and identical distributions of time spent in each state. They differed only in the direction of transitions: the directed chain favored movement around a cycle, whereas the reversible chain had equal probabilities in either direction. Both used the same seven-dimensional state-dependent signal patterns, tested at amplitudes of 1.0 and 0.55, with independent Gaussian noise (SD = 0.60). This allowed us to examine directional transitions separately from state occupancy and signal strength.

We also simulated two continuous processes to test whether HMMs might infer state transitions where none existed. The first generated a shared signal with temporal persistence (autoregressive coefficient 0.90), projected onto 7 networks with fixed weights and independent noise. The second generated a smoothly rotating trajectory in a 2D signal space:

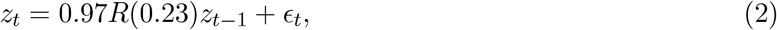

where *R*(0.23) rotates the trajectory by 0.23 radians per time step, and *ϵ*_*t*_ is Gaussian noise (SD = 0.30). Seven projections of this trajectory provided the observed signals, with additional noise (SD = 0.45). Although this process does not switch between discrete states, an HMM might interpret its continuous movement as a sequence of directed transitions.

The four generators, including two amplitudes for each discrete model, gave six conditions. We generated three realizations at each of 12 recording lengths (97–99, 135–140, 159, 197, and 480 frames), giving 216 recordings. We then examined whether changes in how signals were measured could affect model comparisons without altering the underlying dynamics. Each recording was evaluated under four conditions: no filtering, fixed temporal filtering, a change in filtering at the test boundary, and rescaling of the measured signals. Filtering was defined by

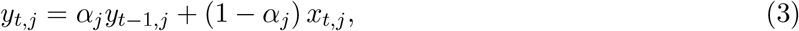

where *x*_*t,j*_ is the generated signal in network j, *y*_*t,j*_ is its filtered value, and *α*_*j*_ controls temporal smoothing. Coefficients ranged from 0.20 to 0.75 across networks, approximating some temporal effects of the hemodynamic response. For the filter-change condition, we reversed the assignment of filter coefficients across networks only during testing, keeping the underlying dynamics and training observations unchanged. This tested whether measurement changes alone could alter the preferred model. Rescaling provided a separate check that comparisons were unaffected by measurement units. The four observation conditions gave 864 cases in total, with matched realizations allowing direct comparisons.

### 2.3 Model Selection

To ensure a fair comparison, we divided each recording into consecutive training (60%), validation (20%), and test (20%) periods. Both models predicted the same held-out observations using preceding time points [15]. The HMM carried its estimated state probabilities across period boundaries, while VAR used preceding measured values. Parameters were fitted on training data, settings selected on validation data, and performance evaluated on test data without refitting. As temporal filtering can affect which model performs better, we allowed each model to use four possible transformations of the observed signals: no transformation, inverse filtering with coefficients of 0.35 or 0.65, and inverse filtering using a separate coefficient for each network, estimated from training data and restricted to the range 0.1–0.8. Inverse filtering attempts to recover the signal before temporal smoothing, using

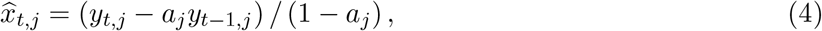

Here *a*_*j*_ is the filter coefficient for network *j*, and the first observation is retained unchanged. We standardized the signals using means and standard deviations from training data only. Prediction densities were converted back to the original measurement units, accounting for the mathematical effects of scaling and inverse filtering. This ensured that the models were compared on the same scale. We tested HMMs with either three or six states, using independent Gaussian observation distributions for each network within a state. Models were initialized using K-means with eight starts, followed by 20 parameter updates. To avoid numerical instability, we imposed a minimum variance of 0.05^2^ and added a total transition pseudocount of 1.5 per row, distributed across destination states. The VAR models predicted each observation from either one or three preceding time points, using ridge regularization (penalty = 1) for the lag coefficients and an unpenalized intercept. Their residual covariance matrices had a minimum of 10^*−*4^*I*. Combining two model sizes with four filtering options gave eight candidates per model family, with all settings fixed throughout the benchmark.

### 2.4 Predictive Density and Directional State Transitions

First, we compared how accurately the models predicted unseen fMRI measurements. For each model, we calculated the average log probability density assigned to the next observed seven-network measurement, given the preceding time points [16]. We then calculated

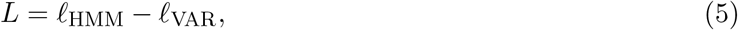

in nats per 7D time point. Positive values indicate better HMM predictions; negative values favor VAR. Second, we tested whether the transitions inferred by the HMM showed a preferred direction; transitions from state A to B might be more frequent than transitions from B to A [17]. We quantified this imbalance using the *stationary transition flux* for fitted transition matrix 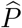 and stationary distribution 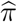:

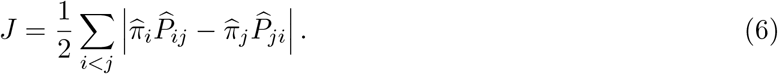

Here 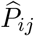 is the probability of moving from state *i* to state *j*, and 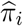 is the long-term probability of occupying state *i*.

The directed chain in Eq. (1) has a flux of 0.07, whereas the reversible chain has zero flux. These measures are designed to answer different questions. Predictive log density assesses how well a model predicts new measurements, while transition flux describes directional differences between its inferred states. Comparing them allows us to test whether directional transitions are specific to discrete dynamics or can also arise when continuous signals are represented by an HMM.

## 3 Synthetic Results

### 3.1 Prediction Across Generating Processes

We evaluated all 864 simulated conditions, fitting 13,824 candidate models. Without temporal filtering, HMMs predicted the data more accurately in 70 of 72 simulations generated by directed state transitions and 69 of 72 generated by reversible transitions. In contrast, VAR performed better in 33 of 36 simulations driven by a shared continuous signal and 34 of 36 simulations with continuous rotational dynamics. Results were similar with fixed temporal filtering: HMMs performed better in 129 of 144 discrete-state simulations, while VAR performed better in 70 of 72 continuous simulations (Table 1). Overall, predictive accuracy generally distinguished the discrete and continuous processes tested here, despite the relatively short recording lengths. In a separate 36-record benchmark, mean HMM transition flux was similar for the discrete cycle (0.068) and continuous rotation (0.057), despite the latter having no discrete states. Flux was lower for the reversible chain (0.018) and common-driver model (0.006). This shows that directional transitions between fitted states can arise from continuous dynamics. Predictive scores from this initial benchmark were not pooled with the main analysis because the test-boundary procedure differed. In the main benchmark with fixed filtering, the directed chain showed greater fitted flux than the reversible chain in 49/60 shorter and 12/12 longer matched pairs using three HMM states, compared with 36/60 and 11/12 using six states. Increasing the number of states therefore did not consistently improve discrimination in short recordings. The six-state HMM was selected in only 151/864 conditions, while first-order VAR was preferred over third-order VAR in every condition.

**Table 1.** Simulations in which HMM predicted held-out signals better than VAR. Columns show no filtering, fixed filtering, or a filter change during testing. Numbers in parentheses give cases per generating process in each column; remaining cases favor VAR.

| World (cases) | Identity | Fixed filter | Test change |
|---|---|---|---|
| Directed (72) | 70 | 66 | 71 |
| Reversible (72) | 69 | 63 | 67 |
| Common AR (36) | 3 | 1 | 34 |
| Rotating AR (36) | 2 | 1 | 34 |

#### Changes in Filtering

Changing only the filter applied during testing increased the number of continuous cases favoring HMM from 2/72 with fixed filtering to 68/72. Each comparison used identical training and validation observations, fitted models, and selected settings; only the test-period filtering changed. Thus, an HMM could appear superior even though the underlying continuous dynamics were unchanged (Fig. 1). Rescaling measurement units preserved all 216 paired density and flux comparisons, apart from negligible numerical errors. Resetting the HMM’s estimated state probabilities at the start of testing rather than carrying them forward, changed mean scores by amounts ranging from *−*0.914 to 0.150 nats/frame. These checks show why model comparisons must account for measurement units and the information available when predictions are made.

**Figure 1.**
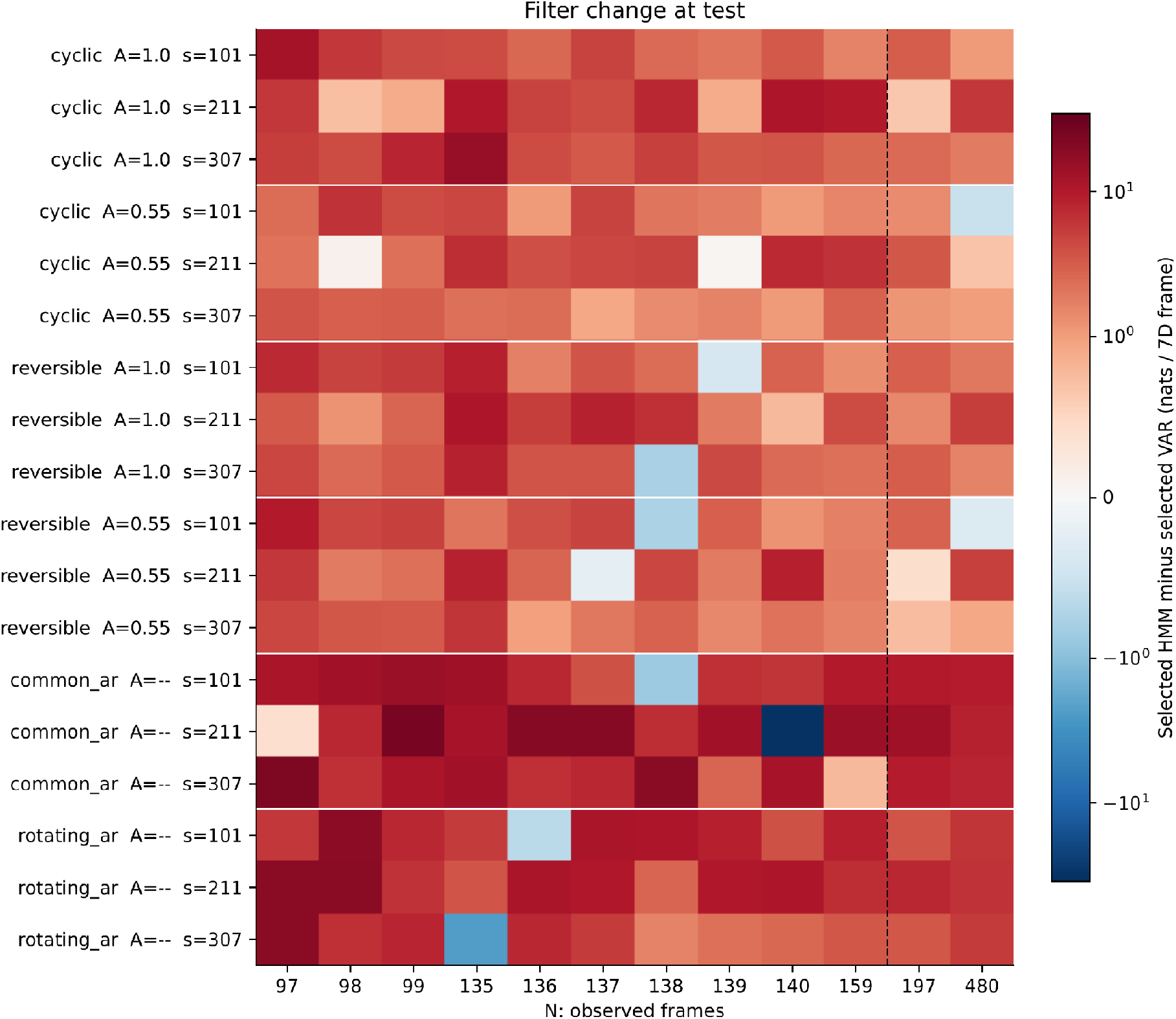
Effect of changing the filter during testing in 216 simulations. Rows show generating process, signal amplitude, and seed; columns show recording length. Red favors HMM and blue favors VAR. Most continuous cases favor HMM despite unchanged underlying dynamics and fitted models. The dashed line separates 197- and 480-frame recordings from shorter ones.

## 4 Human Data Results

We applied the same prediction procedure to the first-visit scans from all 270 ADNI participants, evaluating 4,320 candidate models. VAR predicted the data more accurately in 160/184 recordings with at least 180 frames and 60/86 shorter recordings. The median difference in predictive log density (HMM minus VAR) was *−*1.427 and *−*0.873 nats per seven-network frame, respectively (Fig. 2). All participants were included. The earlier evaluation excluded recordings with fewer than 180 time points, retaining only 184/270 scans. The present analysis included all participants, although estimates of state transitions in shorter recordings remain exploratory. We also tested whether the HMM’s initial state probabilities affected prediction.

**Figure 2.**
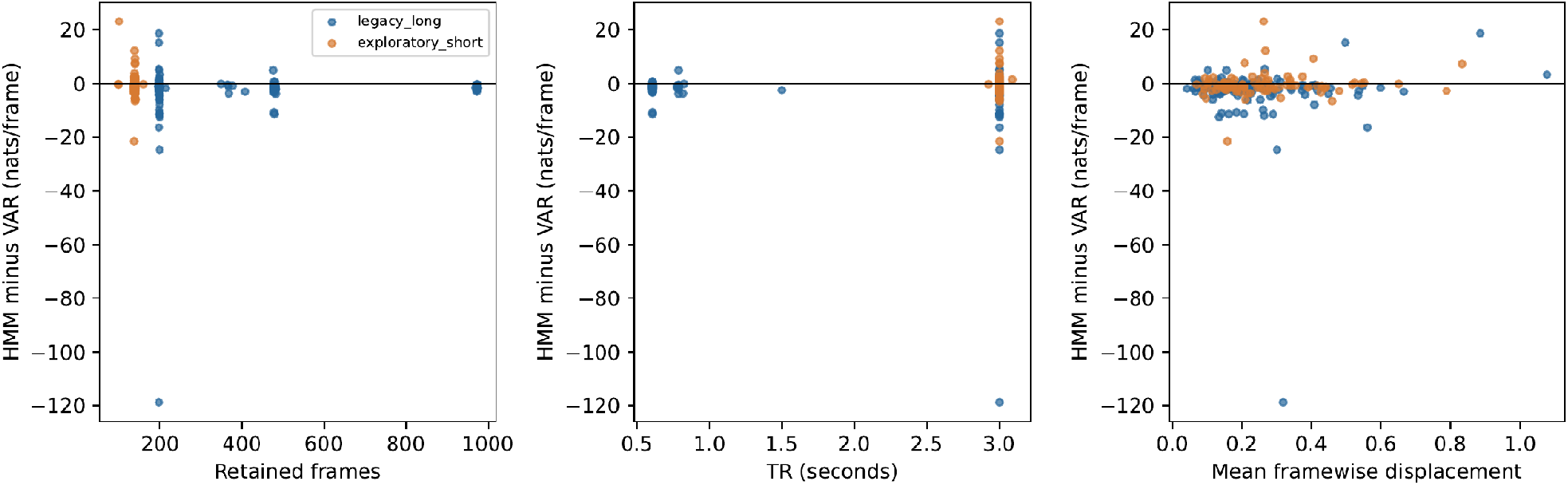
Predictive accuracy of HMM and VAR models in 270 ADNI participants. Each point shows the difference in predictive log density (HMM minus VAR) for one recording, plotted against recording length, repetition time (TR), and mean framewise displacement (a measure of head motion). Positive values favor HMM; negative values favor VAR. Blue indicates recordings with at least 180 time points (n = 184), and orange indicates shorter recordings (n = 86). All participants are shown.

Resetting these probabilities at the test boundary, without changing model parameters, changed the prediction score from *−*0.681 to 0.166 nats/frame.

## 5 Discussion and Limitations

Our results show why transitions between fitted HMM states should not automatically be interpreted as discrete changes in brain activity. Under stable filtering, predictive accuracy generally distinguished the simulated discrete and continuous processes. However, smooth continuous dynamics also produced strongly directional transitions between inferred states. More strikingly, changing only the observation filter reversed the preferred model in most continuous simulations, without changing the underlying dynamics. In ADNI, VAR predicted subsequent fMRI measurements more accurately in most recordings, supporting its use as a benchmark for state-transition models. This does not establish that brain dynamics are continuous rather than discrete. Our findings are limited to the models tested here, particularly HMMs with independent Gaussian observations within states.

More flexible HMMs may perform differently [6, 7]. Short recordings also limit estimates of state transitions [9], while differences in repetition time, head motion, and physiological signals may affect generalization. Overall, interpreting HMM state transitions requires comparison with continuous models and testing whether conclusions remain stable under different observation and preprocessing conditions.

## 6 Code and Data Availability

The generators, complete synthetic observations and all candidate scores are available in the Github repository https://github.com/yury-rd/fMRI-transition.

## 7 Compliance with Ethical Standards

We analyzed previously collected, publicly available ADNI data, without recruiting participants or acquiring new scans. For this analysis of public, de-identified anonymized data, no additional institutional review board approval was required. The synthetic analyses contained no human data.

## Acknowledgments

Data used in preparation of this article were obtained from the Alzheimer’s Disease Neuroimaging Initiative (ADNI) database (adni.loni.usc.edu). As such, the investigators within the ADNI contributed to the design and implementation of ADNI and/or provided data but did not participate in the analysis or writing of this report. A complete listing of ADNI investigators can be found in the ADNI investigator acknowledgment list.

The ADNI imaging and clinical data came from the Alzheimer’s Disease Neuroimaging Initiative database. ADNI began in 2003 as a public-private partnership led by Michael W. Weiner, MD. It brings together longitudinal imaging, biomarkers, and clinical and neuropsychological assessments to study progression from mild cognitive impairment to Alzheimer’s disease.

The National Institute on Aging funds ADNI data collection and sharing through National Institutes of Health grant U19 AG024904 to the Northern California Institute for Research and Education. ADNI has also received support from the National Institute of Biomedical Imaging and Bioengineering, the Canadian Institutes of Health Research, and private contributors through the Foundation for the National Institutes of Health. These contributors include AbbVie, Alzheimer’s Association, Alzheimer’s Drug Discovery Foundation, Araclon Biotech, BioClinica, Inc., Biogen, Bristol-Myers Squibb Company, CereSpir, Inc., Cogstate, Eisai Inc., Elan Pharmaceuticals, Inc., Eli Lilly and Company, EuroImmun, F. Hoffmann-La Roche Ltd and its affiliated company Genentech, Inc., Fujirebio, GE Healthcare, IXICO Ltd., Janssen Alzheimer Immunotherapy Research & Development, LLC., Johnson & Johnson Pharmaceutical Research & Development LLC., Lumosity, Lundbeck, Merck & Co., Inc., Meso Scale Diagnostics, LLC., NeuroRx Research, Neurotrack Technologies, Novartis Pharmaceuticals Corporation, Pfizer Inc., Piramal Imaging, Servier, Takeda Pharmaceutical Company, and Transition Therapeutics.

